# No statistical evidence for an effect of CCR5-∆32 on lifespan in the UK Biobank cohort

**DOI:** 10.1101/787986

**Authors:** Robert Maier, Ali Akbari, Xinzhu Wei, Nick Patterson, Rasmus Nielsen, David Reich

## Abstract

A recent study reported that a 32-base-pair deletion in the *CCR5* gene (CCR5-∆32) is deleterious in the homozygous state in humans. Evidence for this came from a survival analysis in the UK Biobank cohort, and from deviations from Hardy-Weinberg equilibrium at a polymorphism tagging the deletion (rs62625034). Here, we carry out a joint analysis of whole-genome genotyping data and whole-exome sequencing data from the UK Biobank, which reveals that technical artifacts are a more plausible cause for deviations from Hardy-Weinberg equilibrium at this polymorphism. Specifically, we find that individuals homozygous for the deletion in the sequencing data are underrepresented in the genotyping data due to an elevated rate of missing data at rs62625034, possibly because the probe for this SNP overlaps with the ∆32 deletion. Another variant which has a higher concordance with the deletion in the sequencing data shows no associations with mortality. A phenome-wide scan for effects of variants tagging this deletion shows an overall inflation of association p-values, but identifies only one trait at p < 5×10^−8^, and no mediators for an effect on mortality. These analyses show that the original reports of a recessive deleterious effect of CCR5-∆32 are affected by a technical artifact, and that a closer investigation of the same data provides no positive evidence for an effect on lifespan.

## Introduction

CCR5-∆32 is a deletion in the coding region of the *CCR5* gene which has been reported to confer resistance against HIV infections in individuals carrying two copies of the deletion (∆32/∆32)^1,2,3^. Some studies have suggested that the relatively high frequency of this variant in some populations points to a selective advantage conferred by this deletion^4,5^, although the case for natural selection at this variant has also been challenged^6,7^. After the announcement of the birth of two babies whose genomes were edited using CRISPR in order to knock out the *CCR5* gene, additional concerns arose about potential negative effects of this mutation^1,8^.

A recent study by Wei and Nielsen investigated potential deleterious effects in homozygous carriers of this mutation using the UK Biobank data^9^. The study found that a single nucleotide polymorphism (SNP) that tags the CCR5-∆32 deletion (rs62625034) is less common in its homozygous state than expected under Hardy-Weinberg equilibrium (HWE). The study also reported a significantly increased mortality rate in homozygous carriers of this deletion, implying that deleterious effects of this variant, rather than technical artifacts, might be the reason for the deviation from HWE. These findings have been questioned in online discussions, in particular by S. Harrison who focused on whether rs62625034 is indeed a good proxy of the CCR5-∆32 deletion^10^. The recent release of exome sequencing data on around 10% of the UK Biobank samples makes it possible to directly test how well the CCR5-∆32 deletion is tagged by rs62625034 and by other nearby variants. By jointly analyzing the whole genome genotyping data and whole exome sequencing data we find that deviations from HWE in the genotyping data likely are due to technical artifacts. Moreover, when testing for associations of variants in the CCR5-∆32 deletion region to other phenotypes we do not find effects of a magnitude that could explain a strongly increased mortality in ∆32/∆32 individuals.

## Methods

### Markers tagging CCR5-∆32

Genotype data in the UK Biobank is available in three different forms: (1) Allele counts as inferred from the genotyping array intensity values; (2) Imputed genotype dosages which are commonly rounded to best guess allele count integer values. These are based on the array data genotype calls but for many genotyped variants are not equal to the array data genotype calls; and (3) Whole exome sequencing data, currently for a pilot sample of around 10% of the total sample size. Two different pipelines were used to call variants from the read data. Here we only use the variant calls from the GATK pipeline.

We analyse five variants in total, two from the array data, two imputed and one sequenced (Supplementary Table 1):

**rs62625034_genotyped**: This is the genotyped variant which has been used as a proxy for the CCR5-∆32 deletion in Wei and Nielsen^9^.

**rs113010081_genotyped**: A genotyped variant in high LD to the CCR5-∆32 deletion.

**rs113010081_imputed**: The imputed data for the same variant.

**3:46414943_TACAGTCAGTATCAATTCTGGAAGAATTTCCAG_T:** The CCR5-∆32 deletion as called in the imputed data. For brevity, we refer to it as **rs333_imputed**, even though this rs ID is not used in the raw data.

**3:46373452:D:32**: The CCR5-∆32 deletion as called in the exome sequencing data. rs62625034 is not present among the set of imputed SNPs. We refer to it as **rs333_sequenced**, even though this rs ID is not used in the raw data.

It cannot be assumed that any of the analyzed variants perfectly tags the CCR5-∆32 deletion. We treated the direct exome sequencing data on CCR5-∆32 variant itself as the ground truth, and then assess the accuracy of the genotype array variants by comparing them to the exome sequencing variant. We evaluate the concordance between these variants by comparing the counts in each genotype class (0, 1, 2) and computing Pearson correlation coefficients (r^2^). In addition, since the questions of interest relate to the effect of the ∆32/∆32 genotype and we are uninterested for this analysis in misclassification errors between the other two genotypes, we also computed sensitivity and specificity of correctly identifying individuals with two copies of the deletion in the exome sequencing data.

As population heterogeneity can induce deviations from HWE, we limit all of our analyses to individuals classified as “white British” in the UK Biobank. In order to be consistent with Wei and Nielsen^9,11^, we do not exclude related individuals for the results shown here, though our results remain qualitatively the same when excluding related individuals. We compute approximate HWE p-values using a Chi-squared test. However, different data sets have different average deviations from HWE due to the Wahlund-effect and differences in genotype calling data and algorithms. For this reason, Wei and Nielsen^9,11^ used genomic control methods to compute HWE deviations. We follow this protocol^11^ and report two additional sets of HWE p-values. Specifically, we compute HWE p-values by comparing to a set of frequency matched control SNPs (P1), and by using the bootstrap to test for significant deviations from the median value in the genomic control SNPs (P2)^11^. The latter test was argued to be the preferred test as it provides some protection against outliers^11^, which is why it is included here. Each of these methods tests a different null hypothesis, resulting in different p-values.

### Survival rate analysis

To study the effect on survival rates, we extend the phenotypic association analysis by exploring the effects of all variants tagging the CCR5-∆32 deletion on all phenotypes available to us.

For each of the variants, we assess the impact on mortality as previously described in Wei and Nielsen^9,11^. We use five different UK Biobank variables - age at recruitment (ID 21022), Date of attending assessment centre (ID 53), year of birth (ID 34), month of birth (ID 52), and the age at death (ID 40007) - to compute the number of individuals who are ascertained from age i to age i + 1 (N_i_), and the occurrence of death observed from these N_i_ individuals during the interval of age i to age i + 1 (*O*_*i*_). The death rate per year is calculated separately for each ∆32 genotype class as *h*_*i*_ = *O*_*i*_/*N _i_* and the probability of surviving to age i + 1, 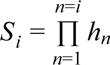. h_77_ is grouped together with h_76_.

To compute p-values for the survival rate analysis, we run Cox proportional hazard models using the ‘coxph’ function in the R-package ‘survival’. We do not use binning into age groups, as described in the previous paragraph, for this analysis. Instead we use only age at recruitment and reported age at death or, if no age at death is reported, the inferred age at time t, where t is the date of the last reported age at death in the entire cohort (16 February 2016).

We estimate power to detect effects on mortality rate in the following way. First, we extract for each sample age at death, or, if age at death has not been reported, the inferred age at time t, where t is the date of the last reported death in the entire cohort (16 February 2016). Next, we randomly draw a genotype (0 or 1) for each person from a Bernoulli distribution with a probability that depends on whether or not this person has died, in proportion to a given relative risk (RR). For individuals who have died, this probability is P(G=1|D) = P(D|G=1) * P(G=1) / P(D), where P(G=1) is the frequency of ∆32/∆32 (0.012), P(D) is the fraction of samples with a reported age at death (0.029), and P(D|G=1) = RR * P(D|G=0) = RR * P(D|G=0) * P(D) / (P(G=1)*P(D|G=1) + (1-P(G=1)) * P(D|G=0)) = RR * P(D) / (P(G=1)*RR + (1-P(G=1))). Similarly, for individuals who are still alive, this probability is P(G=1|A) = P(A|G=1) * P(G=1) / P(A), where P(A) = 1 - P(D) and P(A|G=1) = 1 - P(D|G=1). We then obtain a p-value from a Cox proportional hazard model for each random draw, repeat this 100 times for 9 different RR values, and compute the fraction of random draws with p-value smaller than 0.05 at each value of RR.

### Associations with other phenotypes in the UK Biobank

If a genetic variant has a substantial effect on early mortality then that effect is likely to act through specific phenotypes. We therefore tested whether ∆32/∆32 individuals were at higher risk for 3,331 diseases or disorders than ∆32/+ and +/+ individuals. We tested each of the five variants for associations with 3,911 phenotypes in the UK Biobank. We used the following logistic regression model: y ~ x_01,2_ + c. Here, y is a vector of phenotypes; x_01,2_ is the vector of genotypes, recoded so that each sample with zero or one copy of the deletion is 0 and each sample with two copies of the deletion is 1; and c is a set of covariates, including age, sex, genotyping array, and PC 1 to PC 20, calculated on a set of European individuals^12^. We similarly tested an additional 580 continuous phenotypes using a linear regression model.

We further conducted Poisson tests to check whether any ICD10 diagnosis codes were overrepresented as the reported cause of death in ∆32/∆32 compared to all other individuals.

## Results

### Concordance rates across variants

Figure 1, Supplementary Figures 1 and 2, and Supplementary Tables 2, 3, and 4 show concordance rates, r^2^, and sensitivity and specificity of the genotyped variants and the exome sequencing variant. These analyses suggest that rs113010081_genotyped is a better proxy for the CCR5-∆32 deletion than rs62625034_genotyped.

**Figure 1:**
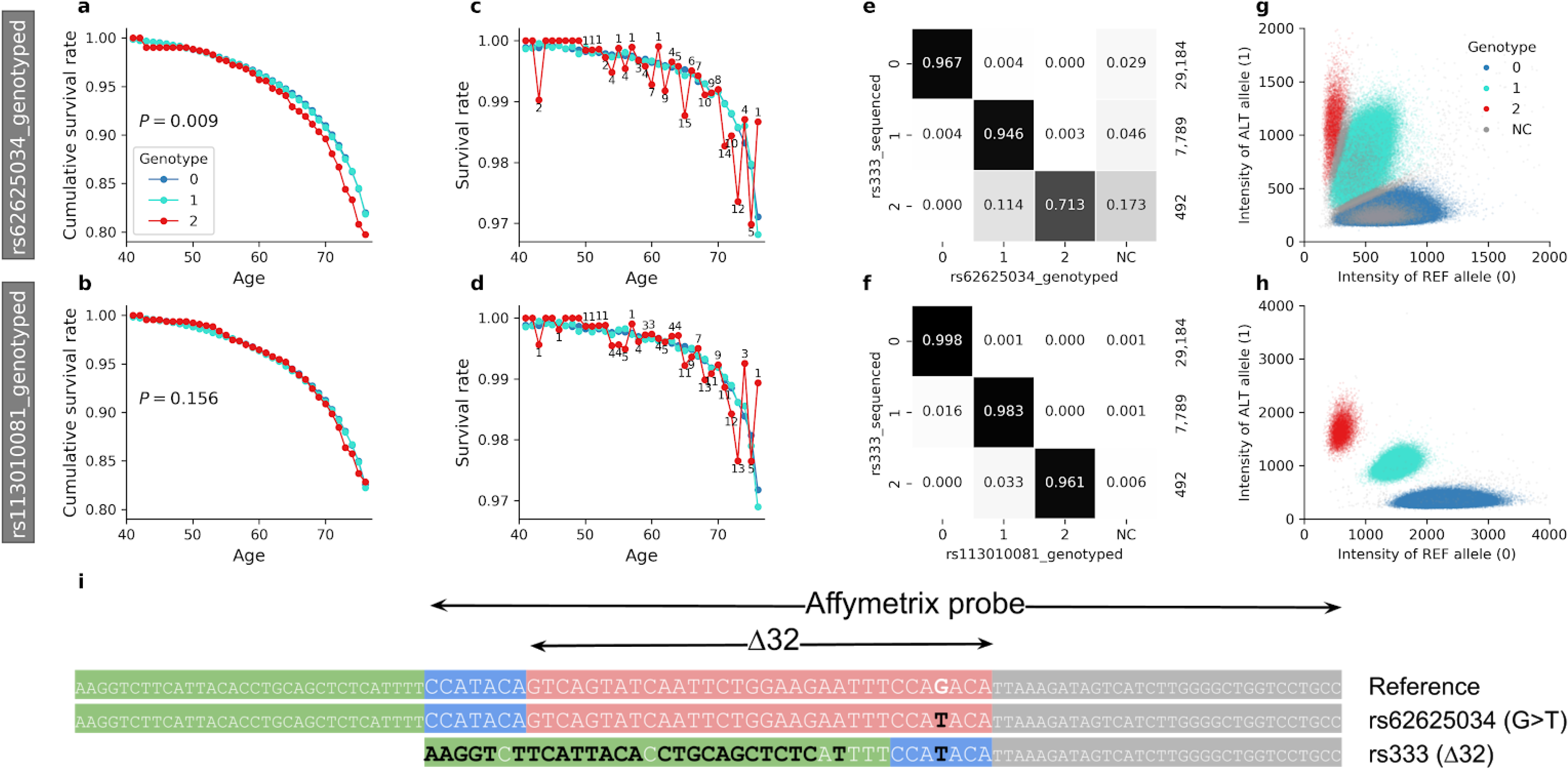
Survival rates for individuals with 0, 1, and 2 copies of the rare allele for two variants tagging the CCR5-∆32 deletion, rs62625034_genotyped (first row) and rs113010081_genotyped (second row). **a, b**, Cumulative survival rates show that the evidence for increased mortality of individuals homozygous for the variant allele in rs62625034_genotyped does not replicate in rs113010081_genotyped. One-sided p-values are from a Cox proportional hazard model which compares survival rates of individuals with 0 or 1 alleles to those with 2 alleles. **c, d**, non-cumulative survival rates, which show the large year-to-year variability in the data caused by small sample counts. Numbers indicate how many ∆32/∆32 individuals have died in each year. **e, f**, Distribution of genotypes at the two variants (including missing genotypes) conditioned on rs333_sequenced genotypes. The total count for each row is on the right. Missing data is strongly correlated to genotype class for rs62625034_genotyped, which fully explains the deviation from Hardy-Weinberg Equilibrium at this site. No such bias is present at rs113010081_genotyped. Numbers are based only on samples genotyped on the UK Biobank Axiom array, as rs113010081_genotyped data is only available for this array **g, h**, Allele intensity clusters for UK Biobank genotyping data, showing the poorer separation of genotype classes for rs62625034_genotyped compared to rs113010081_genotyped. **i**, Different haplotypes at the CCR5-∆32 locus. Black nucleotides differ from the reference. The site of the very rare SNP rs62625034 (G>T) is located within the ∆32 deletion. Due to the sequence similarity at the 3’ end, the probe tags the deletion instead. However, the rs62625034 probes match the reference genotype better than the deletion, leading to higher missingness in the presence of the deletion.

While rs113010081_genotyped has a higher r^2^ with rs333_sequenced than rs62625034_genotyped (0.977 compared to 0.968, Supplementary Table 3), these r^2^ values are mostly influenced by the concordance of the more common genotypes ∆32/+ and +/+. As we are specifically interested in ∆32/∆32 individuals and for the purpose of the present analysis are not concerned by misclassification of the two other much more common genotypes, we also computed sensitivity and specificity based on a comparison of ∆32/∆32 individuals to the union of ∆32/+ and +/+ individuals. The sensitivity and specificity to correctly identify individuals with ∆32/∆32 in the WES data is 0.934 and 0.998 for rs62625034_genotyped, and 0.998 and 1 for rs113010081_genotyped. Out of all individuals identified as ∆32/∆32 by the WES data, 11.4% are classified as ∆32/+ at rs62625034_genotyped, compared to 3.3% at rs113010081_genotyped (Figure 1). This suggests that rs113010081_genotyped more accurately tags ∆32/∆32 individuals than rs62625034_genotyped (Supplementary Table 4).

rs113010081_genotyped was not used by Wei and Nielsen^9,11^ due to its relatively high rate of missing data in the overall dataset. However, detailed examination reveals that the high missingness rate (10.3%) is due to the absence of this variant from the UK BiLEVE Axiom array. This array was used to genotype the first ~ 10% of genotyped samples in the UK Biobank. On the UK Biobank Axiom array, which was used for the remaining ~ 90% of samples, this variant has a missingess rate of 0.08%, while rs62625034_genotyped has a missingness rate of 3.6%, and hence for the individuals for whom we have a genotype at this variant, there are likely to be fewer technical artifacts due to miscalling bias. Indeed, when we examined the probe intensity scatter plots used for genotype calling at the two variants we confirm crisper separation of the genotype classes for rs113010081_genotyped than for rs62625034_genotyped (Figure 1). Supplementary Table 2 shows conditional genotype counts for all individuals, as well as for only those individuals genotyped on the UK Biobank Axiom array. We observed differences in missingness between the two array types, but no relative differences in genotype counts. Other Supplementary Tables only show results from both arrays, as these numbers change very little when restricting to samples genotyped on the UK Biobank Axiom array.

In this work, we do not focus on the imputed variants, as they do not tag the ∆32 deletion as well as the genotyped variants (Supplementary Figure 2 and Supplementary Tables 3 and 4). In addition, Supplementary Table 10 shows that imputation quality differs by genotype at rs11301008_imputed.

### Hardy-Weinberg disequilibrium

We confirm that rs62625034_genotyped shows a highly significant deviation from HWE, caused by a deficiency of individuals with two copies of the rare (deletion tagging) allele (Supplementary Table 5). None of the other tested variants shows a significant deviation from HWE under a Chi-squared HWE test. We tested whether the reduced sample size in the exome sequencing data can explain why we do not see a similarly strong deviation from HWE at the sequenced variant. For this, we compute HWE for all variants also in the subset of samples for which we have exome sequencing data. We find that rs62625034_genotyped still has a HWE p-value of 6.1×10^−9^. In the same subset of samples, the variants rs113010081_genotyped and rs333_sequenced show no deviations from HWE in a Chi-squared test.

To be consistent with Wei and Nielsen, we also compute P1, which measures where the B-statistic (observed/expected ratio of the rare homozygous genotype) falls relative to the distribution of frequency matched control SNPs, and P2, which tests whether the B-statistic of a given variant falls below the median B-statistic of the frequency matched control SNPs (Supplementary Table 5). For rs333_sequenced, the P2 p-value is 0.0276, similar to the previously reported value of 0.0272. For rs62625034_genotyped and rs113010081_genotyped, P2 is < 0.0001 and 0.0023, respectively. P1 is 0.0032 for rs62625034_genotyped, but not significant for the other SNPs. When subsetting to the samples for which we have exome sequencing data, P1 and P2 remain qualitatively similar, however P1 for rs113010081_genotyped is 0.0242.

In comparing inferred genotypes from rs62625034_genotyped and rs333_sequenced, we noticed that 17.3% of individuals who were called as ∆32/∆32 in rs333_sequenced have missing values for rs62625034_genotyped, while only 4.6% and 2.9% are missing for ∆32/+ and +/+, respectively (Figure 1). Correcting for this bias based on the empirically measured differences in missing data rate by genotype class fully explains the discrepancy between sequencing data and genotyping data; that is, the proportion of homozygous minor alleles changes from 1.06% before correction to 1.38% after correction, and the uncorrected HWE p=4.8e-51 becomes p=0.25 after this correction (Supplementary Table 6). In contrast, individuals with missing data at rs113010081_genotyped are not similarly biased with respect to rs333_sequenced (Figure 1).

Figure 1 provides a plausible explanation for why rs62625034_genotyped exhibits higher missingness rates in individuals with the ∆32 deletion. The Affymetrix probe for rs62625034 is targeting a very rare G>T SNP which is located at the 3’ end of the site of the ∆32 deletion. Since this variant is so rare, almost all of the called non-reference alleles indicate the presence of the ∆32 deletion, which at its 3’ end closely resembles the targeted G>T SNP. Since the probe overlaps with the ∆32 deletion but matches it only imperfectly, ∆32 individuals have a higher missingess rate. In contrast, the probe for rs113010081 is 42 kb downstream of ∆32 and suffers from no such problems. In conclusion, rs62625034_genotyped is a biased proxy for ∆32/∆32, while rs113010081_genotyped shows far less evidence of bias.

We carried out a simulation study to test whether increased mortality or other negative ascertainment on ∆32/∆32 individuals can plausibly create a highly significant HWE deviation at this deletion, but no HWE deviation at a SNP with an r^2^ of 0.95 relative to the deletion. We find that ascertainment on one variant induces similarly high deviations from HWE at other variants in high LD (Supplementary Figure 3). Thus, if one variant shows a high degree of deviation from the null expectation of HWE, and another variant in high linkage disequilibrium with it shows no significant deviation from HWE, it is highly likely that a technical artifact is affecting the genotyping of at least one of the variants.

### Survival rate analysis

Confirming the findings of Wei and Nielsen^9,11^, we find that for rs62625034_genotyped, carriers of two copies of the rare allele tend to have a lower survival rate (Figure 1, Supplementary Figure 1, and Supplementary Table 7). We obtain a one-sided p-value from a Cox proportional hazard model of 0.009. However, none of the other tested variants shows any association with survival rate. The fact that the highly correlated rs113010081_genotyped SNP has a p-value of 0.156 when applying the same test, and the small number of deaths per year on which the signal is based (Figure 1) make this finding uncompelling. The power to detect a 20% increased mortality rate at this SNP at a 0.05 significance level is only 75% (Supplementary Figure 4), which means that we cannot rule out that the deletion does affect survival based on this analysis.

Interestingly, we find that samples with missing genotypes at rs113010081_genotyped show greatly increased mortality rates (p-value 2.7×10^−32^). This is a genotyping batch effect: rs113010081_genotyped is absent from the UK BiLEVE Axiom array, and the individuals who were genotyped on this array were ascertained to be smokers^13^. This association disappears when restricting to individuals genotyped on the UK Biobank Axiom array. The same sample restriction does not explain the increased mortality rate seen for two carriers of the rare allele in rs62625034_genotyped (though the p-value increases to 0.016), but this example cautions against reporting associations between variants from the array data and mortality without controlling for possible genotyping array batch effects. We have only observed these batch effects in the array data, but not in the imputed data. Further, we only observed differences in missingness rates between the two array types, but no differences in the relative proportion of called genotypes.

### Associations with other phenotypes in the UK Biobank

CCR5-∆32 is reported to confer HIV resistance only in the presence of two copies of the deletion, and similarly, effects on mortality have also only been reported in the presence of two copies of the deletion. Despite this, association tests for a wide range of phenotypes have only been reported for the additive effect of CCR5-∆32. We therefore tested whether any phenotypes are significantly different in frequency between ∆32/∆32 individuals and all others (∆32/+ and +/+). We tested the five variants for associations with 3,911 phenotypes. Since we think that rs113010081_genotyped most accurately tags ∆32, and the association results for the other variants are similar, we focus on the association results for rs113010081_genotyped. We identify “Lymphocyte count” as the only trait which is significant at a p-value smaller than the classic threshold for declaring genome-wide statistical significance, 5×10^−8^. However, it can be argued that the genome-wide significance threshold is too stringent, since we only test the effect at one locus. When we instead apply Bonferroni multiple testing correction for 3,911 tested phenotypes, we find one additional phenotype, “Mean sphered cell volume”, which is associated at a p-value smaller than 1.27×10^−5^ (which corresponds to 0.05 after Bonferroni correction for 3,911 phenotypes; Supplementary Table 8, Supplementary Figures 5 and 6). The associated phenotypes are similar to the previously reported results from additive association tests and are consistent with CCR5’s role in the immune system. These results suggest that ∆32/∆32 does have effects besides conferring resistance to HIV. We do not observe effects on any diseases which are large enough to explain a substantially increased mortality rate. However, the phenotype “Overall health rating” is associated with rs113010081_genotyped at a nominal p-value of 5.22×10^−3^. On average, ∆32/∆32 individuals are 7% more likely to rate their health as “poor” or “fair” compared to other individuals (Supplementary Table 9). We also obtain p-values of 4.47×10^−3^ and 5.74×10^−3^ for two collections of diagnosis codes described as “Certain infectious and parasitic diseases”^14^. Given that ∆32/∆32 has previously been reported to be a risk factor for symptomatic West Nile virus infection^15^, this is noteworthy. We single out results for these phenotypes because they relate to previously reported effects of ∆32/∆32, but we highlight that we tested almost 4,000 phenotypes. Many other phenotypes with more significant nominal p-values seem unrelated to any relevant health outcomes, and most of these associations are likely due to chance. Despite the large overall sample size, many phenotypes are rare, which further limits the power to detect effects of a genotype present in only 1% of the population at a reasonable significance level.

In analyzing the causes of death, we find no ICD10 codes which are overrepresented in ∆32/∆32 individuals compared to all others, but similar power considerations apply here.

## Discussion

Artificially knocking out a gene in human embryos without fully understanding its function is dangerous, especially as genes that serve no function are unlikely to survive evolutionary pressures. It seems very plausible therefore that there are negative consequences of having no functional copy of the *CCR5* gene. However, our analysis in the UK Biobank of more markers tagging ∆32/∆32 individuals does not provide statistical confirmation that ∆32/∆32 individuals have shorter lifespans than other people. Additionally, in testing associations with a wide range of phenotypes we do find weak effects on a handful of traits, but none that suggest a 20% increased mortality rate.

Deviations from HWE can be due to causal genetic effects on mortality or on other phenotypes which lead to survival biases. However, technical artifacts are a much more common cause for deviations from HWE. Our analysis suggests that the rs113010081_genotyped variant tags the ∆32 better than rs62625034_genotyped, without showing any evidence of HWE deviation. The HWE deviation observed at this SNP is, therefore, almost certainly caused by a higher missingess rate at ∆32/∆32 individuals. Thus, deviations from HWE at rs62625034_genotyped cannot be interpreted as evidence for a deleterious effect of the ∆32/∆32 genotype.

Beyond the specific explorations into the phenotypic effects of the CCR5-∆32/∆32 genotype, this study highlights the delicacy of association analysis. Specifically, it provides a case example of the subtle pitfalls that can produce false-positive results, even in an extraordinarily high quality and relatively uniformly generated dataset like the UK Biobank. Our re-examination of previously published results was inspired by S. Harrison’s identification of qualitatively inconsistent findings between two variants in strong linkage disequilibrium with each other^10^ which we replicated and further explored here.

## Acknowledgements

This research has been conducted using the UK Biobank Resource under Application Number 31063. We acknowledge the participants in the UK Biobank. We are grateful to Benjamin Neale and Alkes Price for critical comments, and to Sean Harrison for a blog posting that showed how the association results and HWE p-values at rs113010081_genotyped were qualitatively discordant with those at rs62625034_genotyped, which prompted us to re-examine these issues. We thank Konrad Karczewski, Kári Stefansson and Mark Daly for sharing with us early versions of two other manuscripts re-examining the evidence of association to mortality at *CCR5*, and working with us to post all manuscripts together. This work was funded in part by NIH grants GM100233 and HG006399, the John Templeton Foundation grant 61220, and the Howard Hughes Medical Institute.

## Author contributions

R.M., A.A. performed analyses all except for the one presented in Supplementary Table 10, which was carried out by X.W. N.P., R.N., and D.R. supervised the study. R.M., A.A., and D.R. wrote the manuscript with critical review from all co-authors.

---

**Supplementary Table 1:**
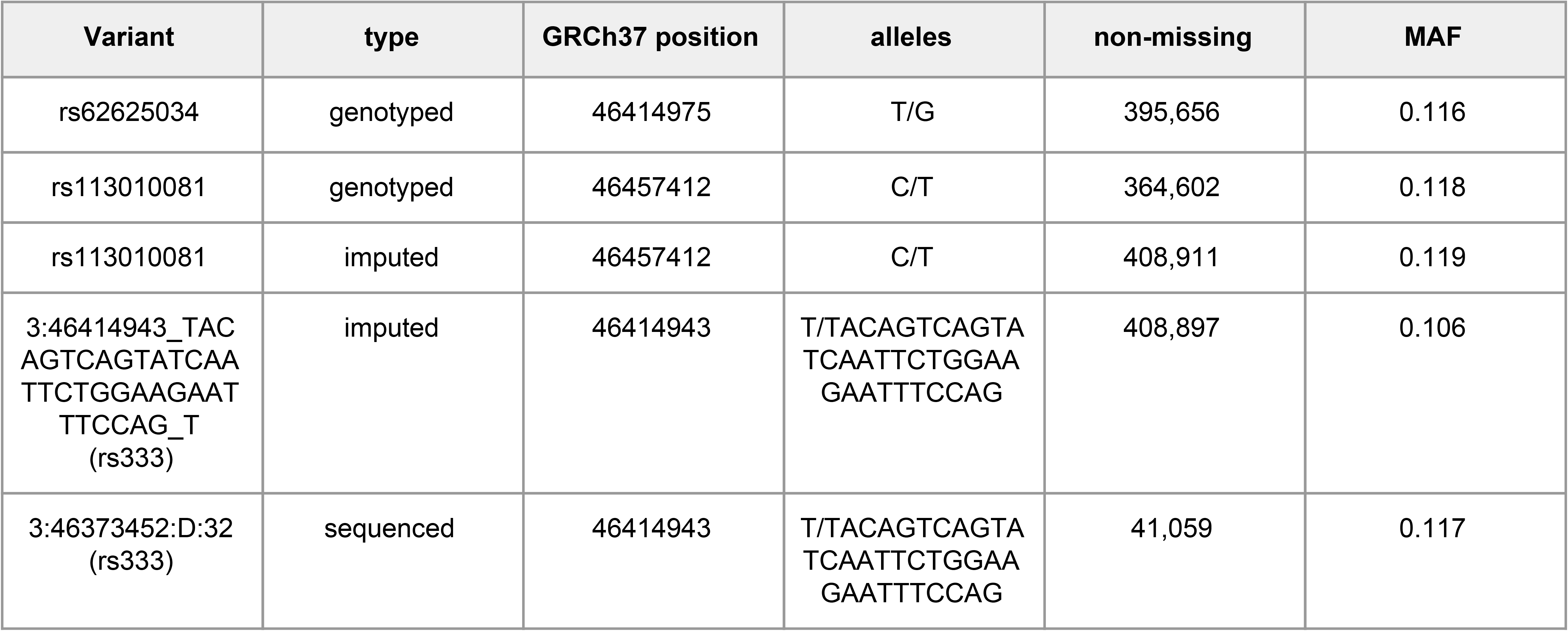
Variants tagging the CCR5-∆32 deletion.

**Supplementary Table 2:** Cross-tabulation of allele counts for genotyped variants tagging the CCR5-∆32 deletion against the variant in the exome sequenced data.

| Variant | Allele count<br>non-sequenced | Allele count sequenced<br>(rs333_sequenced)<br>All samples |  |  | Allele count sequenced<br>(rs333_sequenced)<br>Only UK Biobank Axiom array |  |  |
| --- | --- | --- | --- | --- | --- | --- | --- |
|  |  | 0 | 1 | 2 | 0 | 1 | 2 |
| rs62625034_genotyped | 0 | 30984 | 35 | 0 | 28212 | 33 | 0 |
| rs62625034_genotyped | 1 | 129 | 8094 | 66 | 120 | 7370 | 56 |
| rs62625034_genotyped | 2 | 0 | 27 | 380 | 0 | 25 | 351 |
| rs62625034_genotyped | NA | 872 | 381 | 91 | 852 | 361 | 85 |
| rs113010081_genotyped | 0 | 29127 | 126 | 0 | 29127 | 126 | 0 |
| rs113010081_genotyped | 1 | 34 | 7658 | 16 | 34 | 7658 | 16 |
| rs113010081_genotyped | 2 | 0 | 1 | 473 | 0 | 1 | 473 |
| rs113010081_genotyped | NA | 2824 | 752 | 48 | 23 | 4 | 3 |
| rs113010081_imputed | 0 | 31671 | 279 | 0 | 28901 | 257 | 0 |
| rs113010081_imputed | 1 | 246 | 8200 | 37 | 217 | 7480 | 34 |
| rs113010081_imputed | 2 | NA | 42 | 500 | 0 | 37 | 458 |
| rs113010081_imputed | NA | 68 | 16 | 0 | 66 | 15 | 0 |
| rs333_imputed | 0 | 31696 | 1150 | 5 | 28918 | 1032 | 5 |
| rs333_imputed | 1 | 221 | 7347 | 156 | 200 | 6720 | 146 |
| rs333_imputed | 2 | 0 | 24 | 375 | 0 | 22 | 340 |
| rs333_imputed | NA | 68 | 16 | 1 | 66 | 15 | 1 |

**Supplementary Table 3:** r^2^ between variants the CCR5-∆32 deletion.

| Variant | rs333_sequenced | rs62625034_genotyped | rs113010081_genotyped | rs113010081_imputed |
| --- | --- | --- | --- | --- |
| rs62625034_genotyped | 0.968 |  |  |  |
| rs113010081_genotyped | 0.977 | 0.946 |  |  |
| rs113010081_imputed | 0.930 | 0.900 | 0.950 |  |
| rs333_imputed | 0.818 | 0.789 | 0.808 | 0.803 |

**Supplementary Table 4:** Sensitivity and specificity of genotyped variants to correctly identify samples with two copies of the CCR5-∆32 deletion in the exome sequencing data.

| Variant | P | N | TP | TN | Sensitivity (TP/P) | Specificity (TN/N) |
| --- | --- | --- | --- | --- | --- | --- |
| rs62625034_genotyped | 407 | 39308 | 380 | 39242 | 0.934 | 0.998 |
| rs113010081_genotyped | 474 | 36961 | 473 | 36945 | 0.998 | 1.000 |
| rs113010081_imputed | 542 | 40433 | 500 | 40396 | 0.923 | 0.999 |
| rs333_imputed | 399 | 40575 | 375 | 40414 | 0.940 | 0.996 |

**Supplementary Table 5:** HWE p-values for variants tagging the CCR5-∆32 deletion. In brackets: observed and expected number of samples with two copies of the rare allele.

| Variant | Chi-squared HWE p-values |  | P1 and P2 p-values (genomic control corrected) |  |
| --- | --- | --- | --- | --- |
|  | All samples | Samples with WES data | All samples | Samples with WES data |
| rs333_sequenced | 0.22 (537, 562) | 0.22 (537, 562) | N/A | 0.0764, 0.0276 |
| rs62625034_genotyped | 4.8e-51 (4348, 5317) | 6.1e-09 (421, 540) | 0.0032, < 0.0001 | 0.0022, < 0.0001 |
| rs113010081_genotyped | 0.36 (4979, 5036) | 0.23 (496, 520) | 0.0941, 0.0023 | 0.0242, 0.0326 |
| rs113010081_imputed | 0.78 (5759, 5778) | 0.48 (565, 580) | N/A | N/A |
| rs333_imputed | 1.4e-05 (4301, 4563) | 0.02 (416, 461) | N/A | N/A |

**Supplementary Table 6:**
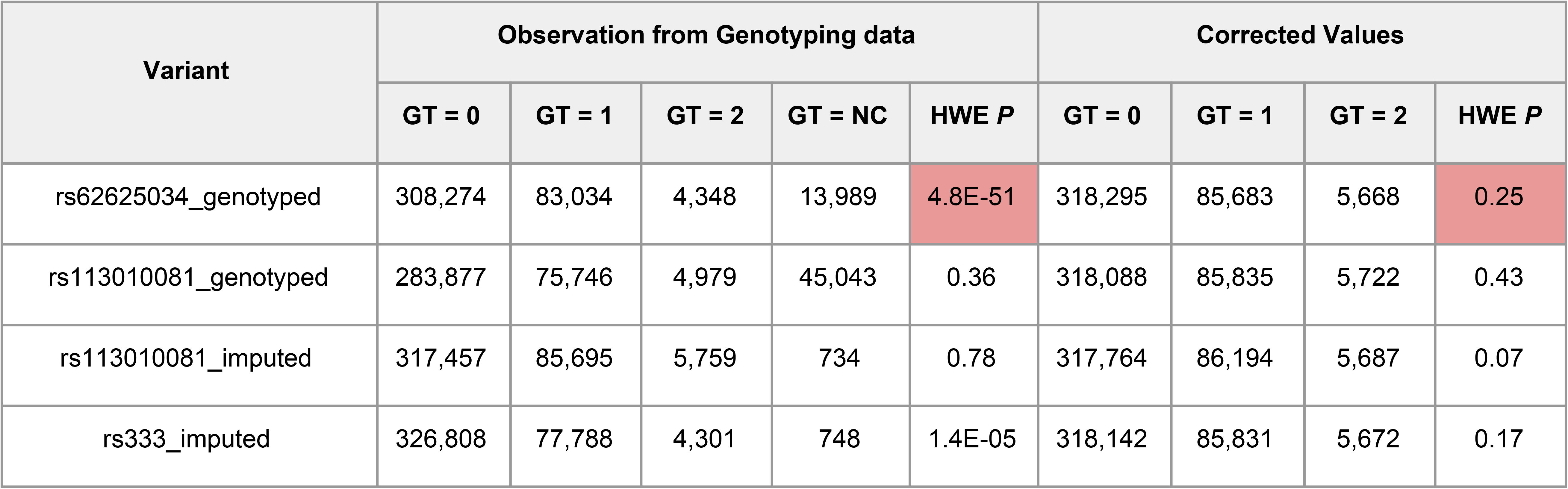
Correcting for bias can explain the extreme p-value for the violation of HWE for rs62625034_genotyped. Unbiased genotype counts is the expected number of true genotypes conditioned on observations in the genotyping array data (including missing genotypes). Conditional distribution is estimated by the joint distribution of genotyping array and UK Biobank WES data. UK Biobank WES data is considered as the ground truth. This table includes all white British samples in the UK Biobank.

**Supplementary Table 7:** Number of samples who have died, for each variant and age group. Values correspond to the red dots in the third row of Supplementary Figure 1.

| Variant | 41 | 42 | 43 | 44 | 45 | 46 | 47 | 48 | 49 | 50 | 51 | 52 | 53 | 54 | 55 | 56 | 57 | 58 | 59 | 60 | 61 | 62 | 63 | 64 | 65 | 66 | 67 | 68 | 69 | 70 | 71 | 72 | 73 | 74 | 75 | 76 |
| --- | --- | --- | --- | --- | --- | --- | --- | --- | --- | --- | --- | --- | --- | --- | --- | --- | --- | --- | --- | --- | --- | --- | --- | --- | --- | --- | --- | --- | --- | --- | --- | --- | --- | --- | --- | --- |
| rs333_sequenced | 0 | 0 | 0 | 0 | 0 | 0 | 0 | 0 | 0 | 0 | 0 | 0 | 0 | 1 | 1 | 2 | 0 | 1 | 0 | 0 | 0 | 0 | 0 | 0 | 2 | 0 | 1 | 2 | 0 | 0 | 0 | 1 | 1 | 0 | 1 | 0 |
| rs62625034_genotyped | 0 | 0 | 2 | 0 | 0 | 0 | 0 | 0 | 0 | 1 | 1 | 1 | 2 | 4 | 1 | 4 | 1 | 3 | 4 | 7 | 1 | 9 | 4 | 5 | 15 | 6 | 7 | 10 | 9 | 8 | 14 | 10 | 12 | 4 | 5 | 1 |
| rs113010081_genotyped | 0 | 0 | 1 | 0 | 0 | 1 | 0 | 0 | 0 | 1 | 1 | 1 | 1 | 4 | 4 | 5 | 1 | 4 | 3 | 3 | 4 | 5 | 4 | 4 | 11 | 9 | 7 | 13 | 11 | 9 | 11 | 12 | 13 | 3 | 5 | 1 |
| rs113010081_imputed | 0 | 0 | 2 | 0 | 0 | 1 | 0 | 0 | 1 | 1 | 1 | 1 | 1 | 4 | 3 | 6 | 1 | 4 | 3 | 7 | 4 | 6 | 4 | 5 | 16 | 9 | 11 | 14 | 11 | 16 | 14 | 13 | 13 | 4 | 6 | 1 |
| rs333_imputed | 0 | 0 | 1 | 0 | 0 | 0 | 0 | 0 | 1 | 1 | 0 | 1 | 1 | 3 | 3 | 5 | 2 | 2 | 3 | 7 | 3 | 3 | 4 | 4 | 9 | 9 | 5 | 10 | 10 | 10 | 10 | 10 | 10 | 5 | 6 | 1 |

**Supplementary Table 8:**
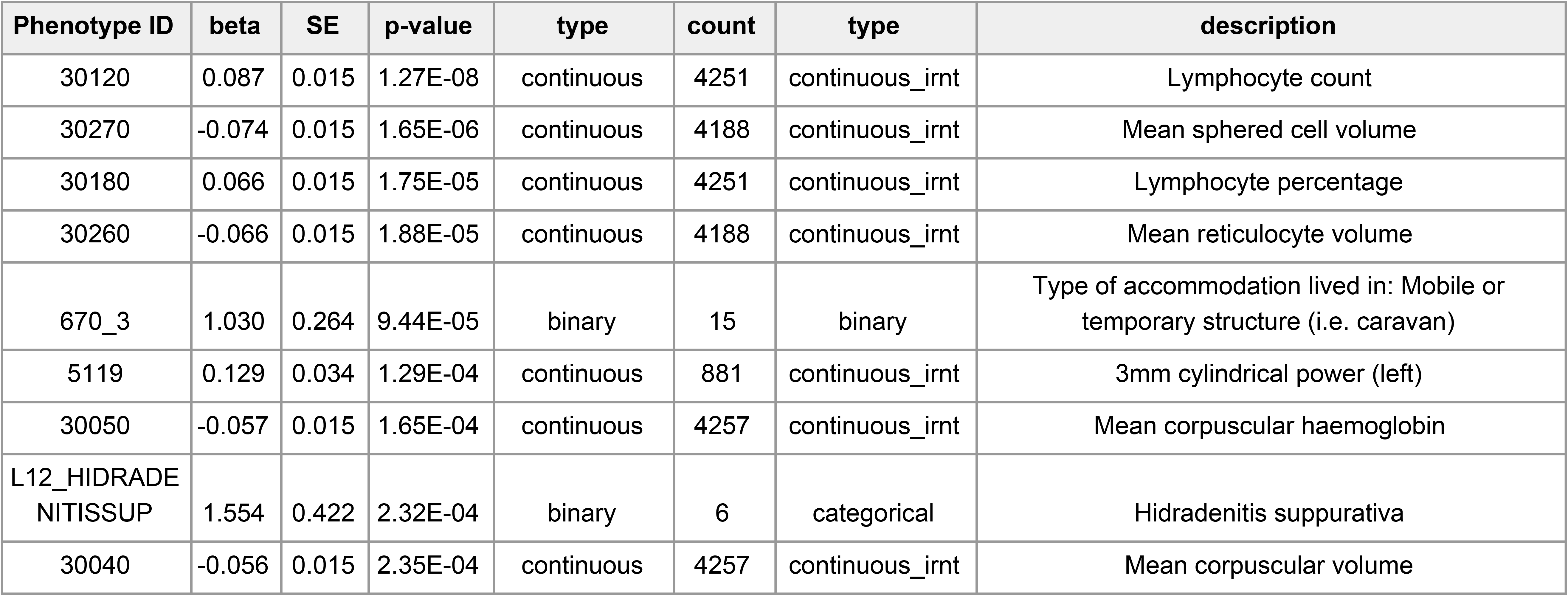
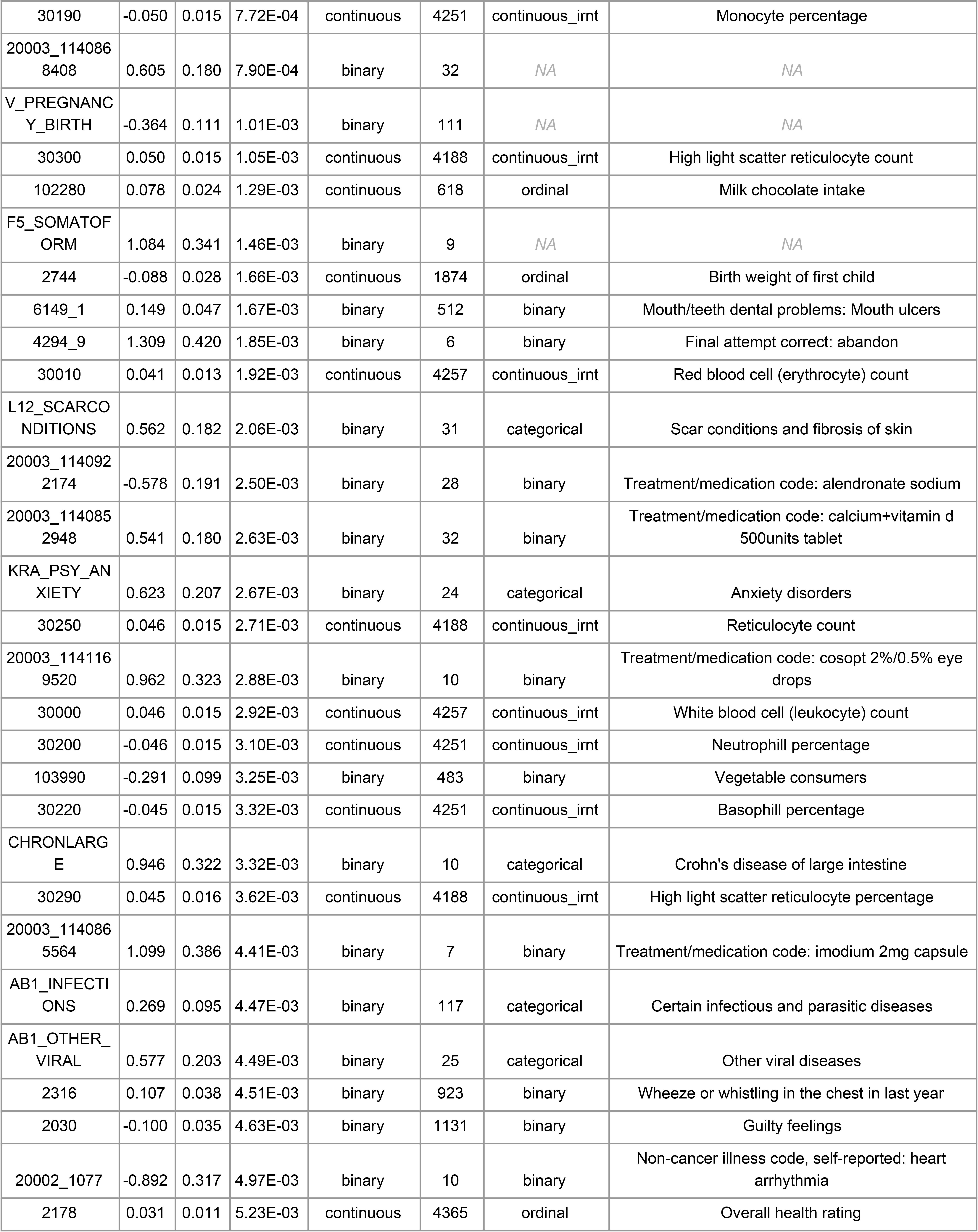
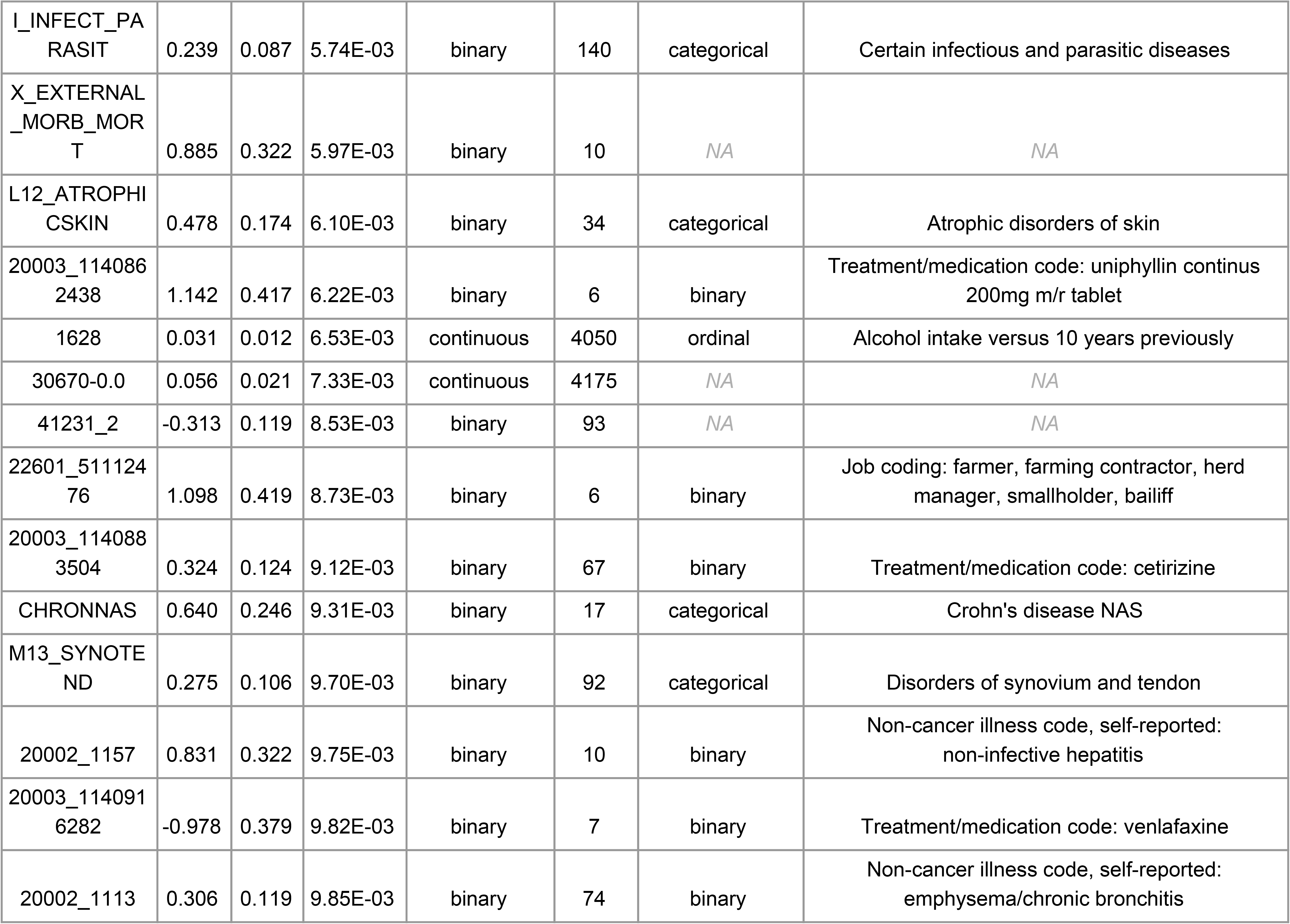
Association results for rs113010081_genotyped showing phenotypes with p-value < 0.01. Phenotypes with p-value < 1.27×10^−5^ are significant after Bonferroni correction for 3,911 phenotypes. The count column lists the number of ∆32/∆32 individuals who are cases (for binary phenotypes) or who have non-missing phenotype information (for all other phenotypes).

**Supplementary Table 9:** Contingency table of self-reported health rating and ∆32 status inferred from rs113010081_genotyped. The odds ratio of “Fair” or “Poor” health vs “Excellent” or ”Good” health is 1.068. Adjusted and unadjusted p-values are 0.0052 and 1.

| Overall health rating | $\Delta 32/+$ and $+/+$ | $\Delta 32/\Delta 32$ observed | $\Delta 32/\Delta 32$ expected |
| --- | --- | --- | --- |
| Excellent | 54436 | 676 | 751.59 |
| Good | 185795 | 2592 | 2569.13 |
| Fair | 62757 | 913 | 868.30 |
| Poor | 12720 | 184 | 175.98 |

**Supplementary Table 10:**
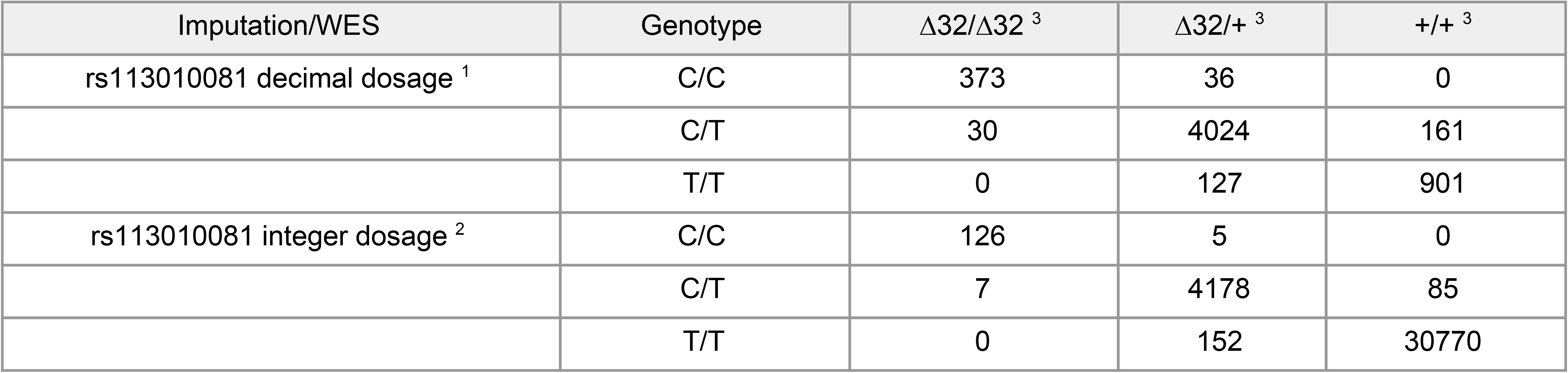
Genotype calls at rs11301008_imputed and rs333_sequenced in the UK Biobank White British. ^1^Individuals with imputed dosage (0,0.5] as C/C, (0.5,1.5) as C/T, and [1.5,2) as T/T. ^2^ Individuals with imputed dosage 0 as C/C, 1 as C/T, and 2 as T/T. ^3^Individuals with ∆32/∆32, ∆32/+, and +/+ genotypes in the UK Biobank WES data. Notice the relative increase in ∆32/∆32, ∆32/+ genotypes with decimal dosage (low confidence imputation) relative to integer dosage (high confidence imputation), and the relative large discrepancy between the exome sequencing data and imputation based genotyping data for decimal dosage genotypes. For example, within the class of genotypes with decimal dosage, 30/403 homozygous minor genotypes in the exome sequencing data are called as heterozygous in the UK Biobank decimal dosage imputation data, and 36/409 homozygous minor genotypes in the UK Biobank decimal dosage imputation data are called as heterozygous in the exome sequencing data.

| Imputation/WES | Genotype | $\Delta 32/\Delta 32$ <sup>3</sup> | $\Delta 32/+$ <sup>3</sup> | $+/+$ <sup>3</sup> |
| --- | --- | --- | --- | --- |
| rs113010081 decimal dosage <sup>1</sup> | C/C | 373 | 36 | 0 |
|  | C/T | 30 | 4024 | 161 |
|  | T/T | 0 | 127 | 901 |
| rs113010081 integer dosage <sup>2</sup> | C/C | 126 | 5 | 0 |
|  | C/T | 7 | 4178 | 85 |
|  | T/T | 0 | 152 | 30770 |

**Supplementary Figure 1:**
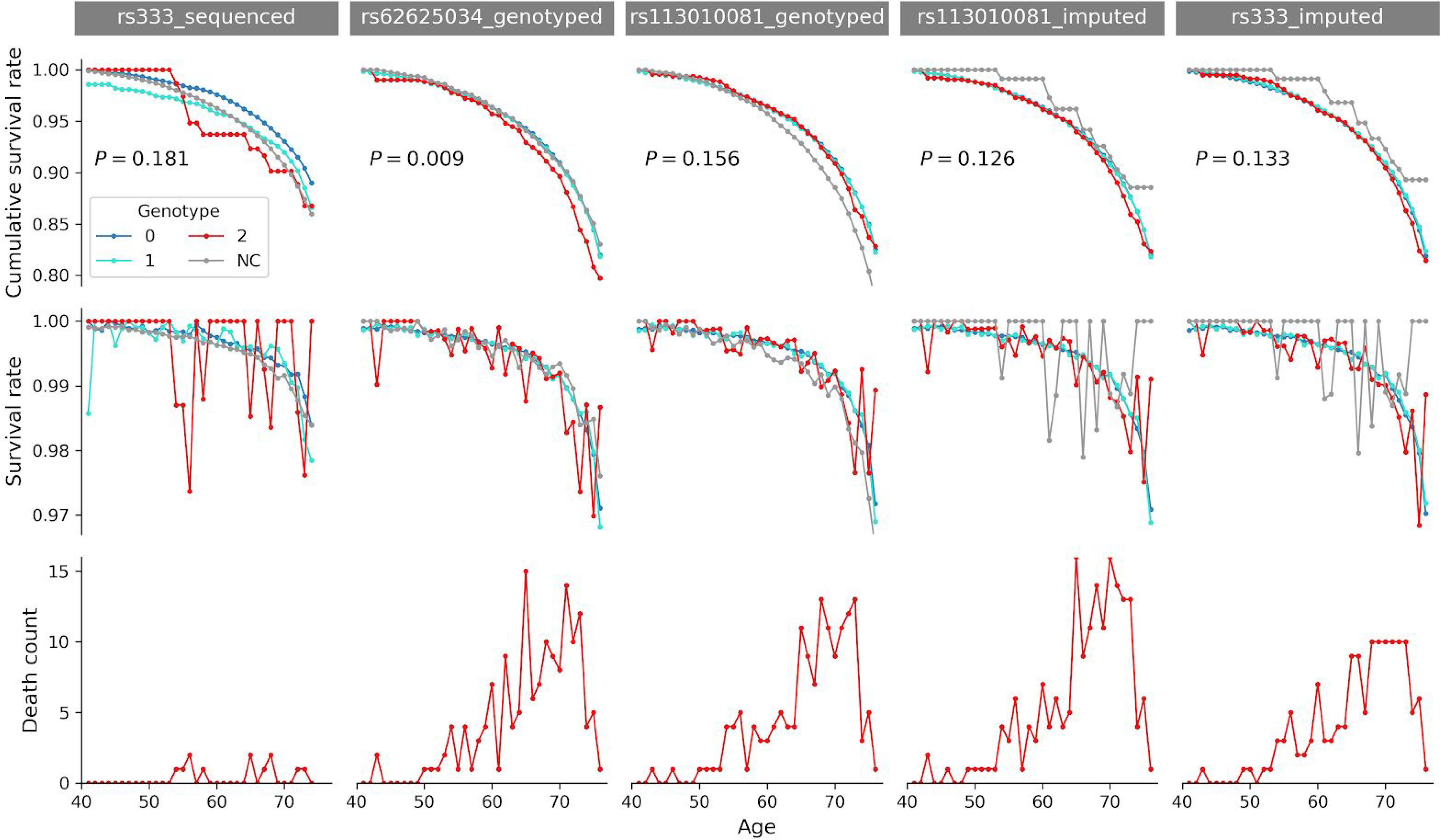
Survival rates for individuals with 0, 1, or 2 copies of the rare allele or No Call (NC) for variants tagging the CCR5-∆32 deletion. First row: Cumulative survival rates. Numbers are one-sided p-values of a Cox proportional hazard model which compares survival rates of individuals with 0 or 1 alleles to those with 2 alleles. Second row: non-cumulative survival rates. Third row: Number of individuals who have died in any given year with 2 copies of rare allele (see also Supplementary Table 7).

**Supplementary Figure 2:**
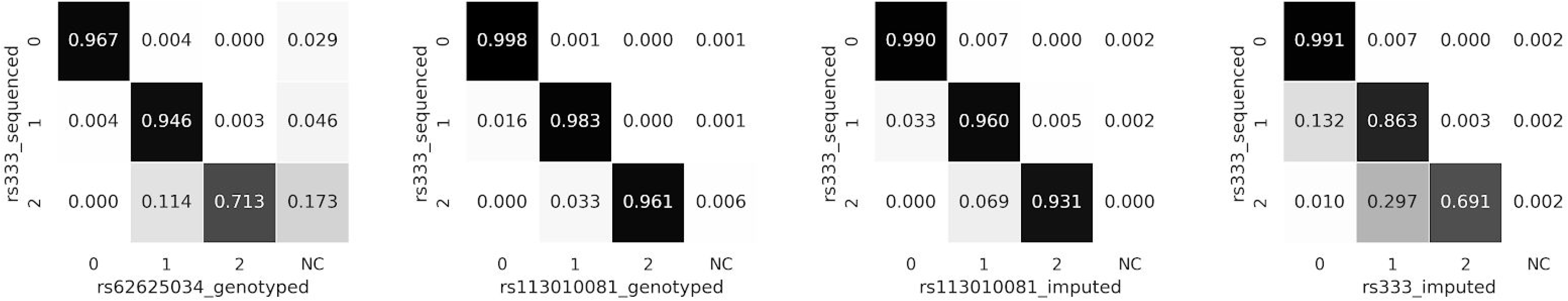
Confusion matrix for different markers with missing data. The last column of the first panel shows that individuals with missing genotype at rs62625034_genotyped are enriched for ∆32/∆32 according to rs333_sequenced. This can lead to a violation of HWE at rs62625034_genotyped. All white British samples of UK Biobank WES data shared with UK Biobank Axiom array data are used in this figure.

**Supplementary Figure 3:**
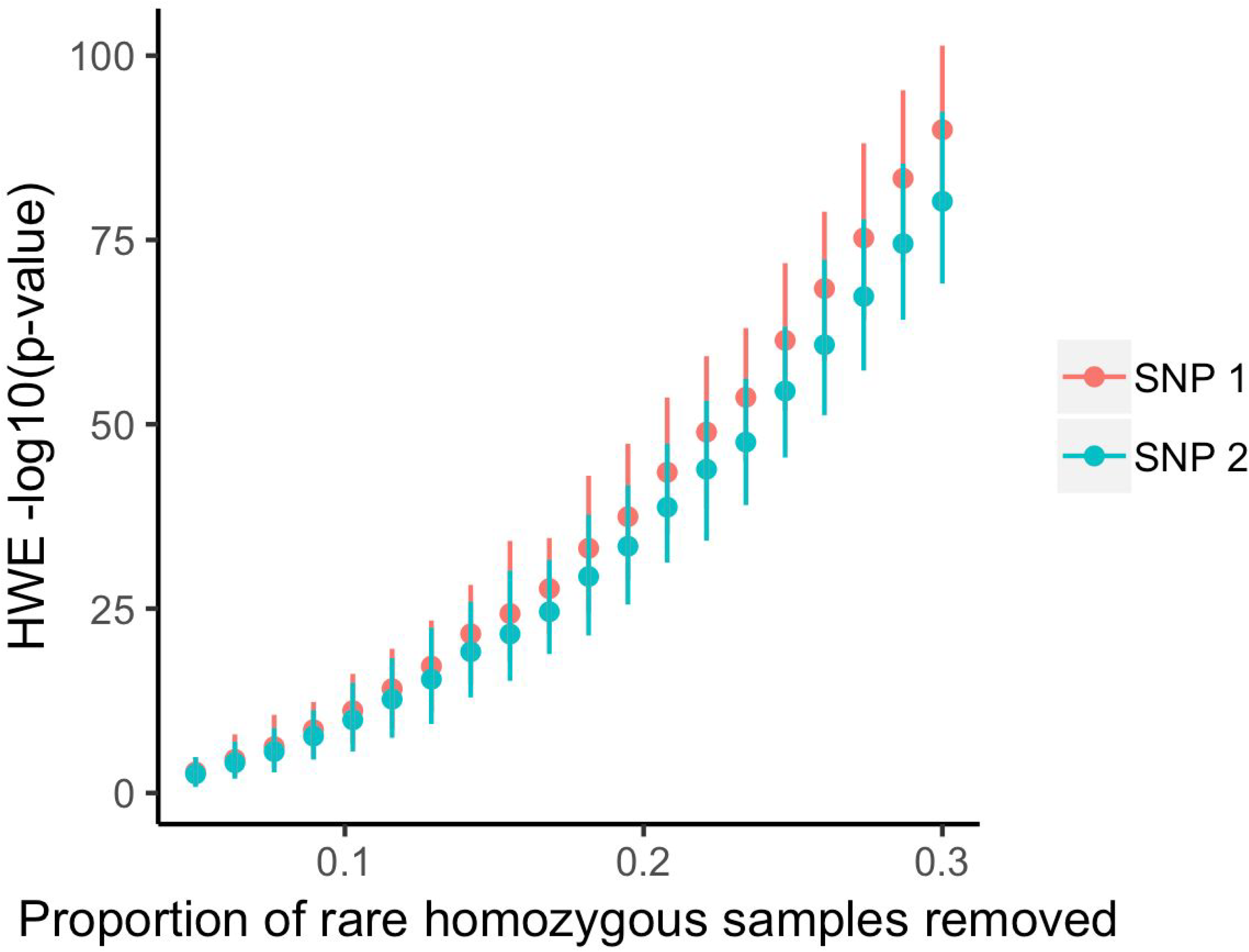
Simulated HWE Chi-squared p-values at two variants with minor allele frequency of 11% with r^2^ of 0.95, in a sample of 400,000 individuals. Both variants are initially in HWE. We then remove a subset of samples which are homozygous for the rare allele at SNP 1. This leads to a deviation from HWE at SNP 1, but it also leads to a similar deviation from HWE at SNP 2. Only simultaneous selection acting in the opposing direction on SNP 2, or technical artifacts which create a dependence of missingness in one SNP on genotype in the other SNP explain a situation where HWE p-values are very different at both SNPs. Error bars denote the 5th and 95th percentile out of 100 replicates in each bin.

**Supplementary Figure 4:**
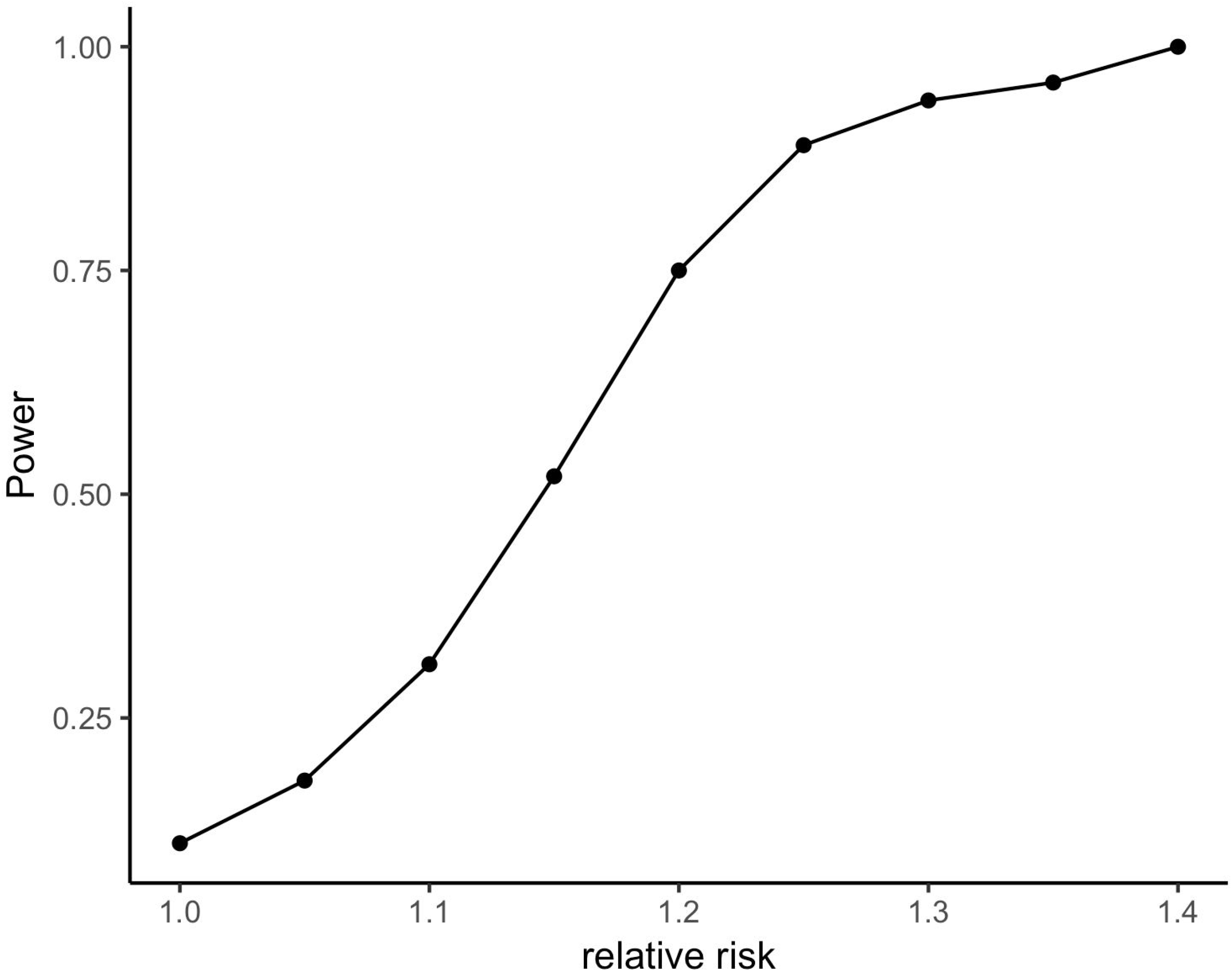
Power to detect effects on mortality of a genotype with the frequency of ∆32/∆32 in a sample of the same total size and mortality rate as the cohort studied here, as a function of relative risk. The power to detect a 20% increase in mortality rate at a 0.05 significance level is 75%.

**Supplementary Figure 5:**
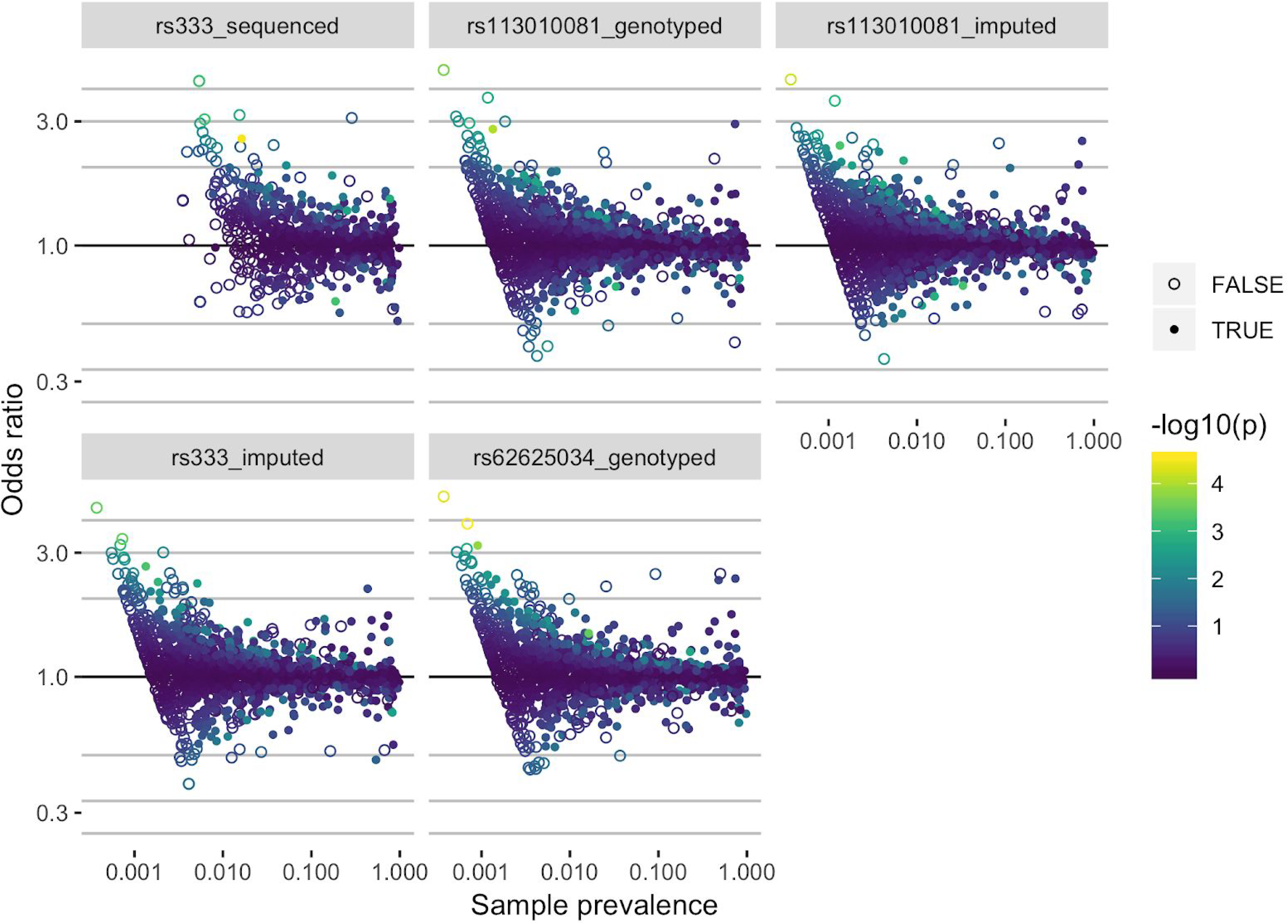
Odds ratios (*e*^β^) for all case-control phenotypes in five variants as a function of sample prevalence. Colors represent uncorrected p-values. Open circles represent case-control phenotypes with 10 or fewer cases in ∆32/∆32 individuals. Only phenotypes with more than five cases in ∆32/∆32 individuals are shown.

**Supplementary Figure 6:**
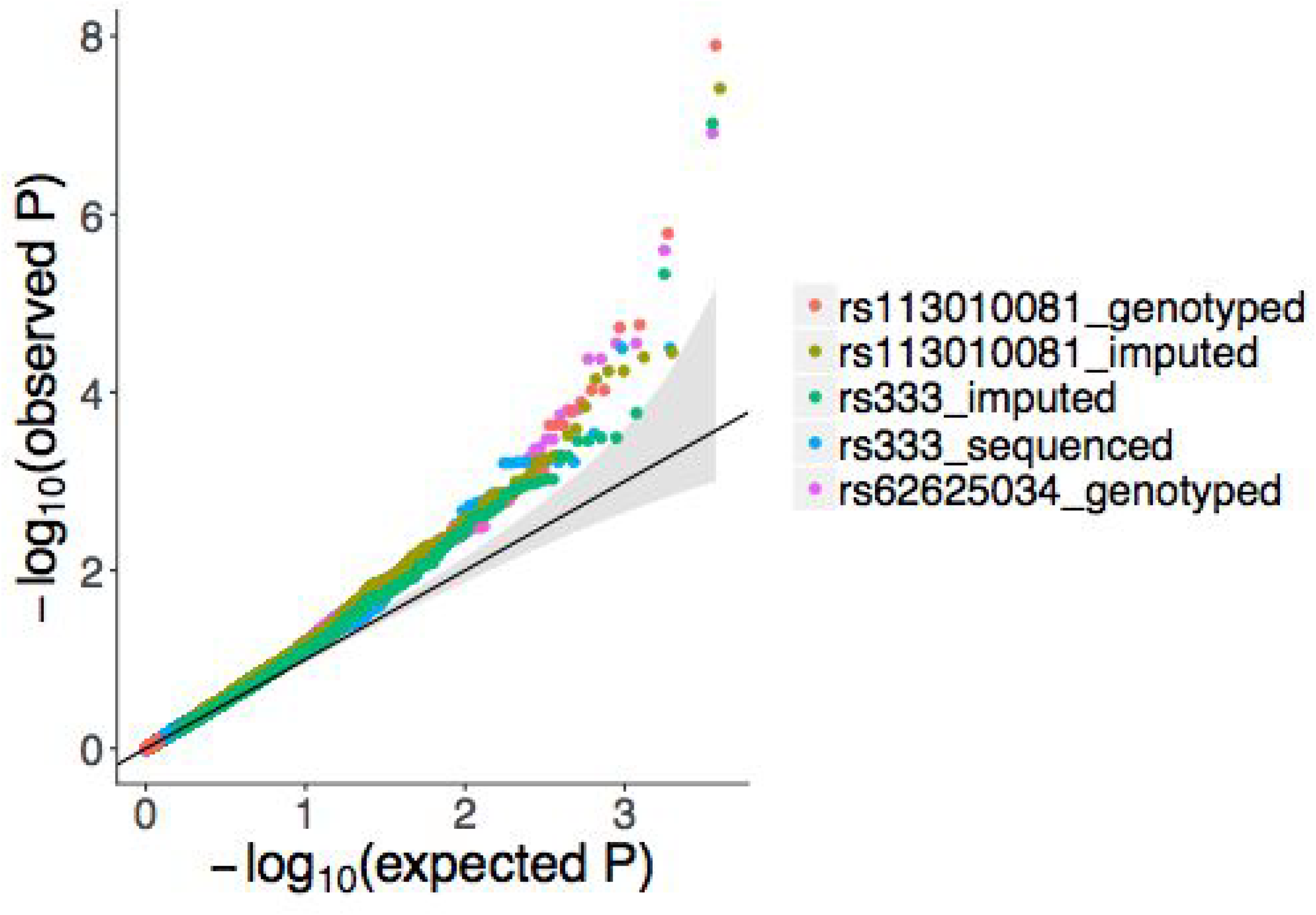
QQ-plot of the associations across all phenotypes. Each variant is plotted in a different color. Only phenotypes with more than five cases in ∆32/∆32 individuals are shown.

